# Screening of surface exposed lipoproteins of *Leptospira* involved in modulation of host innate immune response

**DOI:** 10.1101/2021.08.20.457056

**Authors:** Ajay Kumar, Vivek P. Varma, Syed M. Faisal

## Abstract

*Leptospira*, a zoonotic pathogen is capable of causing both chronic and acute infection in susceptible host. Surface exposed lipoproteins play major role in modulating the host immune response by activating the innate cells like macrophages and DCs or evading complement attack and killing by phagocytes like neutrophils to favour pathogenesis and establish infection. In this study we screened some of surface exposed lipoproteins which are known to be involved in pathogenesis for their possible role in immune modulation (innate immune activation or evasion). Surface proteins of Len family (LenB, LenD, LenE), Lsa30, Loa22 and Lipl21 were purified in recombinant form and then tested for their ability to activate macrophages of different host (mouse, human and bovine). These proteins were tested for binding with complement regulators (FH, C4BP), host protease (plasminogen, PLG) and as nucleases to access their possible role in innate immune evasion. Our results show that of various proteins tested Loa22 induced strong innate activation and Lsa30 was least stimulatory as evident from production of pro-inflammatory cytokines (IL-6, TNF-a) and expression of surface markers (CD80, CD86, MHCII). All the tested proteins were able to bind to FH, C4BP and PLG, however Loa22 showed strong binding to PLG correlating to plasmin activity. All the proteins except Loa22 showed nuclease activity albeit with requirement of different metal ions. The nuclease activity of these proteins correlated to in vitro degradation of Neutrophil extracellular trap (NET). These results indicate that these surface proteins are involved in innate immune modulation and may play critical role in assisting the bacteria to invade and colonize the host tissue for persistent infection.

## Introduction

Leptospirosis is a zoonotic disease caused by spirochetes bacteria which belong to genus *Leptospira*. With 1.03 million cases and 58,900 deaths per year it has become leading cause of morbidity and mortality especially in tropical countries(Ko et al., 2009). Lack of early detection and unavailability of vaccines capable of inducing cross protection against various serovars are major hurdles in containment of infection and reducing associated pathology. Little is known about Leptospira pathogenesis and host immune response which has hampered development of effective vaccine(Ko et al., 2009). Pathogenic bacteria including *Leptospira* upon entering a susceptible host is recognized by innate immune system which tries to kill the bacteria by variety of mechanism. Pattern recognition receptors like TLRs and NLRs upon recognition induces activation of innate immune cells and subsequent induction of adaptive response(Santecchia et al., 2020) Complement system comprising of soluble proteins in blood initiate cascade of reaction and leading to rapid killing of pathogen by phagocytosis and target cell lysis(Dunkelberger and Song, 2010) Phagocytes like Neutrophils may kill the bacteria by generating reactive oxygen species (ROS), cytotoxic granules, antimicrobial peptides and generating Neutrophil Extracellular Traps (NETs)(Teng et al., 2017). To escape from these innate defences pathogen may induce TLR dependent inflammatory response to favour their pathogenesis but may also escape recognition through this protective response by antigenic variation or downregulating the expression of surface proteins. It is likely that *Leptospira* evades recognition from TLR2 and TLR4 possibly by modulating the expression of surface proteins as mice lacking these receptors were highly susceptible to infection(Chassin et al., 2009). *Leptospira* may acquire complement regulators and host proteases to escape from complement mediated killing. To avoid killing by phagocytes like neutrophils they exploit their surface proteins to evade extravagation and chemotaxis, opsonisation and phagocytosis(Teng et al., 2017).. They may express surface proteins as nucleases that can degrade NET once it gets entrapped as has been reported in other pathogens (Thammavongsa et al., 2013;Doke et al., 2017;Storisteanu et al., 2017).

Previous studies identified several surface proteins of *Leptospira* as pro-inflammatory and capable of inducing TLR2 or TLR4 dependent activation of innate immune response(Hung et al., 2006;Yang et al., 2006;Wang et al., 2012;Faisal et al., 2016) Several surface proteins have also been shown to bind FH, C4BP and PLG to evade from complement mediated killing (Fraga et al., 2016). LipL21 has shown to modulate neutrophil function by inhibiting MPO(Vieira et al., 2018). Recently, *Leptospira* has been shown to induce formation of NET, however expression of any nuclease that can degrade NET has not been reported yet(Scharrig et al., 2015). Although several surface proteins of *Leptospira* have been identified which are involved in activation or evasion from host innate immune response, however till date no attempt is made to identify those proteins which are involved in immune modulation (both activation and evasion). Based on literature review, we selected some of the surface exposed lipoproteins like LenB, LenD, LenE, Lsa30, Loa22 and LipL21 known to be involved in pathogenesis to access their role in immune modulation(Raja and Natarajaseenivasan, 2015). *Leptospira* endostatin like proteins (Len proteins) are family of outer membrane lipoprotein comprising Len A, Len B, Len C, Len D, Len E, LenF which are widely distributed in pathogenic serovars. They have important role in pathogenesis as they are expressed during infection and have been shown to interact with components of host extracellular matrix like laminin and fibronectin(Vieira et al., 2014). Len A and Len B have been shown to bind to FH and FHR-1(Verma et al., 2006;Stevenson et al., 2007). Loa22 is surface protein having a OmpA like domain which is conserved in pathogenic serovars (Ristow et al., 2007). Loa22 is expressed during infection and is strongly recognized by sera obtained from human and bovine leptospirosis (Gamberini et al., 2005;Ghosh et al., 2018). It has shown to bind to components of ECM like fibronectin and collagen and has shown to mediate proinflammatory response possibly through TLR2 recognition(Barbosa and Isaac, 2020;Hsu et al., 2021) (Barbosa et al., 2006). It is an essential virulence factor as Loa22 mutant was attenuated in virulence(Ristow et al., 2007). LipL21 is second most abundant surface protein after LipL32 and is conserved among pathogenic serovars(Cullen et al., 2003). It is immunogenic and being recognized by both immune sera from human and hamsters infected with *Leptospira (Cullen et al., 2003*;*Ratet et al., 2017)*. Lsa30 is novel adhesin which is shown to bind with ECM components, interact with C4BP and plasminogen(Souza et al., 2012;Fouts et al., 2016).

In the present study, we screened these proteins to assess their possible role in innate immune modulation i.e. ability to activate macrophages, bind to complement regulators and host proteases and act as nucleases for their possible role in degradation of NET. We cloned, expressed and purified these proteins in recombinant form and tested their ability to activate mouse, human and bovine macrophages. We tested the binding ability of these proteins with complement regulators (FH, C4BP) and host proteases (PLG). Further we tested the nuclease activity of these proteins and correlated this nuclease activity with degradation of NET in vitro.

## Material & Methods

### Cell lines and reagents

A mouse, bovine ad human macrophage cell lines RAW264.7, BoMac and THP-1 respectively were originally purchased from the American Type Culture Collection (Manassas, VA) USA. Cells were cultured in specified medium (DMEM or RPMI, Sigma, USA) supplemented with 10% FBS (Invitrogen, Carlsbad, CA, USA), penicillin (100 U/ml), and streptomycin (100 mg/ml) and maintained at 37°C, 5% CO_2_. IL-6 and TNF-α cytokine sandwich ELISA kits specific for human or mouse or bovine were purchased from R&D biosystems. PE-CY5– conjugated anti MHC-II, PE-conjugated CD86, APC conjugated CD80, for flow cytometry experiments were procured from BD Biosciences, US. Complement reagent Serum (Sigma S1-100ml), Goat anti FH, Rabbit anti C4BP, Mouse anti PLG (Sigma SAB1406263-50UG), Plasmin substrate, UPA (SIGMA SRP6273).

### Cloning, expression and purification of recombinant proteins

Low-passage virulent *Leptospira interrogans* serovar Pomona was cultured at 28°C in Ellinghausen–McCullough–Johnson–Harris (EMJH) medium (BD Difco™, USA) supplemented with 10% bovine serum albumin (BSA). Genomic DNA was isolated using kit. Genes coding for surface exposed lipoproteins viz. LenB, LenD, LenE, Lsa30, Lipl21 and Loa22 were PCR amplified using specific primer and successfully cloned into His tagged Pet28a SUMO vector. Sequence of the cloned gene was verified with T7 primers. The plasmid constructs were transformed into BL21 DE3 and resulting transformants were grown at 37 °C overnight as primary culture and secondary culture in 1 L of LB broth containing kanamycin (50 μg/ml); the expression of the protein was induced with 1 mM isopropyl β-D-1-thiogalactoside (IPTG). The cells were harvested by centrifugation at 3,000 rpm and the cell pellet was resuspended in 100 mM Tris Cl, 150 mM Nacl pH8.0 and followed by sonication at constant pulses. The lysate was centrifuged to remove cell debris and the supernatant was subjected to affinity chromatography using Ni-NTA beads (GE Life Sciences). The recombinant fusion proteins were eluted in 300 mM Imidazole in PBS pH 7.4while the protein was bound to the column by incubating at 4 °C for 12 to 16 h. The proteins were eluted and checked for size and purity by SDS-PAGE. The concentration of purified protein was estimated using Bradford reagent (Thermo Fischer).

### Cell stimulation assays

RAW264.7 cells (1 × 10^5^ cells/well) were cultured in complete DMEM medium and THP-1 cells (1 × 10^5^ cells/well) were cultured in complete RPMI medium and differentiated to macrophage in presence of 100nm phorbol 12-myristate 12-acetate (PMA) for 3 days at 37°C and 5%CO2. Thereafter, PMA was removed and fresh medium was added. BoMac cells (1 × 10^5^ cells/well) were cultured in complete RPMI medium. Cells were stimulated with varying concentration (1, 2, 5ug/ml) of each protein rLenB, rLenD, rLenE, rLsa30, rLipl21 and rLoa22 for 24hrs at 37°C in presence of 5%CO_2._ Cells stimulated with PAM3CSK4 (20ng/ml,200ng/ml,) and LPS (500ng/ml,100ng/ml,) were used as positive controls whereas those stimulated with media alone were considered as negative control. The proteins were pre-treated with Polymyxin B (10ug/ml protein) at 37°C for 1hr or Proteinase K (5μg/ml protein) at 65°C for 1hr followed by inactivation at 95°C for 5min before each assay to rule out endotoxin activity.

### Cytokine ELISA

The culture supernatant from stimulated RAW264.7 and THP-1 cells were collected after 24hrs and cytokines (IL-6, TNF-a) were measured using sandwich ELISA kits (R & D systems) following manufacturers protocol.

### RT-PCR

After 4 h of treatment BoMac cells were recovered in 500μl of TRIzol (Invitrogen, Carlsbad, CA) and equal volumes of chloroform were added; samples were centrifuged at 12000rpm for 15min at 4°C. The aqueous phase was then passed through RNA easy mini columns (MN) and RNA was purified following the manufacturer’s protocol. The RNA quantity was assessed by UV spectroscopy and purity by 260/280 ratio. First-strand cDNA was synthesized using the Takara following the manufacturer’s instructions. RT-PCR was performed in 96 well microtiter plates in a 10μl reaction volume containing 50ng cDNA, 10μM each primer Table-1 and SYBR green (Bio Rad). Samples were run in triplicate and data was analysed with Sequence Detection System (Bio-Rad CFX-96). The experimental data were presented as fold changes of IL-6 and TNF-a gene expression. RNA levels of the analyzed genes were normalized to the amount of GAPDH present in each sample.

### Flow cytometry analysis

RAW264.7 (10^6^ cells/well) were stimulated with PAM3CSK4(20ng/ml) or LPS(500ng/ml) or PMB treated (2ug/ml) rLenB or rLenD or rLenE or rLsa30 or rLipl21 or rLoa22 for 24 h at 37°C in presence of 5%CO_2_. Cells were harvested and washed with prechilled PBS and after blocking they were then incubated with PE-CY5–conjugated anti MHC-II, PE-conjugated CD80, PE vio conjugated CD86, (BD biosciences, USA) for 1 h on ice in the dark. The cells were fixed with 1% paraformaldehyde and 50,000 total events/sample were acquired using a FACScan calibrator. The data were analysed using FlowJo software.

### Dot Blot Binding Assay

Dot blot binding assays were performed to confirm the binding of various recombinant proteins rLenB, rLenD, rLenE, rLsa30, rLipl21 and rLoa22 with complement regulators FH/C4BP/PLG. Protein 1ug/ml were immobilized onto NC membranes (0.22-m pore size; Bio-Rad). The membrane kept for drying for 5-10 min. The membranes were blocked with 5% BSA in Trisbuffered saline–Tween 20 (TBS-T) for 2hrs at RT, washed three times with TBS-T, and incubated with 10% NHS diluted in PBS with gentle shaking for 3hr at RT. After extensive washing with TBS-T, the blots were incubated with the corresponding primary antibody (anti-FH, anti-C4BP, anti-PLG; 1: 5,000 dilution) in TBS-T for 2 hrs at RT. The blots were washed with TBS-T and incubated with a peroxidase-conjugated anti-goat, anti-rabbit and anti-mouse secondary antibody (1: 6,000 or 1: 6,000 dilution) for 2 hrs at room temperature. Reactive spots were developed using a chemiluminescence system.

### ELISA binding assay

Protein attachment to soluble complement regulators FH and C4BP, and PLG was analysed by ELISA. Micro titre plates were coated overnight at 4^0^C with 1 µg/ml each rLenB, rLenD, rLenE, rLsa30, rLipl21 and rLoa22. BSA was used as negative control. The wells were washed three times with PBS containing 0.05 % Tween 20 (PBS-T), blocked with 300ul PBS/2 % BSA for 2 h at 37 C, and incubated with different concentration (0–100 %) of normal human serum (NHS) diluted in PBS for 90 min at 37^0^C. After washing, binding between recombinant protein and each component was detected by adding appropriately diluted rabbit anti-C4BP or goat anti-factor H or mouse anti-PLG antibodies and further incubated for 1 h at 37^0^C. After washing the plates (three times), 100 µl PBS containing horseradish peroxidase (HRP)-conjugated anti-rabbit/goat/mouse IgG was added to each well and plates were incubated for 1 h at 37^0^C. After usual washing, TMB (100 µl per well) was added and reactions were allowed to proceed for 15 min and stopped by the addition of 50 ml 2N H_2_SO_4_. The plate was read at 450 nm in a microplate reader.

### Plasmin activity assay

Microtitre plate wells were coated overnight with 2µg/well rLenB, rLenD, rLenE, rLsa30, rLipl21, rLoa22 at 4 C overnight. The wells were washed with PBS-T and blocked for 2 h at 37^0^C with 10% non-fat milk. The blocking solution was discarded, and human PLG (2ug/well) was added, followed by an incubation for 90 min at 37^0^C. Wells were washed three times with PBS-T, and then 3U per well uPA was added together with plasmin-specific substrate D-Val-Leu-Lys 4-nitroanilide dihydrochloridein and plate was incubated for 24h, 48h and 72h at 37^0^C. The absorbance was measured at 405nm.

### Nuclease activity

To examine the DNase activity of of rLenB, rLenD, rLenE, rLsa30, rLipl21 and rLoa22, DNA fragment was incubated with 5µg concentration of each protein or DNase I (20IU, positive control) in DPBS with 5mM MgCl_2_ or CaCl_2_ or ZnCl_2_ in a PCR tube at 37°C for 2h. The reaction mixture was subjected to EtBr Agarose gel electrophoresis (1%) and then observed under the Gel doc. Cleavage activity was quantified by measuring the ethidium bromide signal in each lane and calculating the fraction of DNA digested relative to the untreated DNA.

### Isolation of Neutrophils from murine blood

Neutrophils were isolated from mouse blood using standard procedure. Blood was collected from 4-5 C57BL/6 mice by ocular puncture in tubes containing EDTA. Blood (1ml) pooled from different mice was mixed with ACK lysis buffer to lyse RBCs. The cells were washed with RPMI 1640 supplemented with 10% FBS, counted and re-suspended in 1ml of ice-cold sterile PBS. Cells were overlaid on 3ml of Histopaque 1077/1119 mix in a 15ml conical tube and then centrifuged for 30min at 825*g at 25°C without braking. Neutrophils at the interface were collected and washed twice with a complete RPMI-1640 medium, counted, and suspended in the medium for the specific assay. The viability was determined by Trypan blue exclusion assay. The purity of neutrophil was determined by FACS using anti-Ly6G antibody.

### NET assay

NETosis was measured by quantification of elastase released from degraded NET DNA using the Neutrophil Elastase Activity Assay kit (Cayman, Chemical) following manufacturers protocol. Briefly, mouse neutrophils (1 × 10^5^ cells/well) were seeded in 96-well plate and treated with 3µl of PMA (50ng/ml) followed by incubation for 3h at 37°C/5%CO2. After NET induction, the medium was replaced with 5ug protein (rLenB or rLenD or rLenE or rLsa30 or rLipl21 or rLoa22) in nuclease assay buffer (PBS with 5 mM MgCl2 or CaCl2 or ZnCl2) and incubated for 4 hrs at 37°C /5%CO2. DNaseI and BSA was used as a positive and negative control respectively. At the end of treatment,10 ul of supernatant was mixed with 90 ul diluted assay buffer and finally 10 ul substrate solution (Z-Ala-Ala-Ala-Ala) was added. The total mixture was incubated in black OPTI plate for 1.5 hrs at 37°C. The plates were read in multi-mode reader at an excitation wavelength of 485 nm and an emission wavelength of 525 nm.

### Statistical Analysis

For all the experiments, wherever required, GraphPad Prism 7.0 (GraphPad Software, Inc.) and one-way ANOVA were executed for the analysis of the results. The data were represented as the mean of triplicates ± SEM. p < 0.05 was considered significant.

## Results

### Cloning, expression, and purification of recombinant proteins

The surface exposed lipoproteins involved in pathogenesis viz. Len proteins (LenB, LenD,LenE), Lsa30, Loa22 and LipL21 were selected based previous studies as compiled in a review article by V Raja et. al (Raja and Natarajaseenivasan, 2015). The genes coding for LenB, LenD, LenE, Lsa30, Lipl21 and Loa22 were PCR amplified, cloned, expressed and purified from soluble fractions. The SDS-PAGE profile in Fig. 1 shows that proteins were highly pure with expected molecular weight of LenB (27kDa), LenD (61.94kDa), LenE (65KdA), Lsa30 (45kDa), Lipl21 (21kDa) and Loa22 (22kDa).

**Fig. 1:**
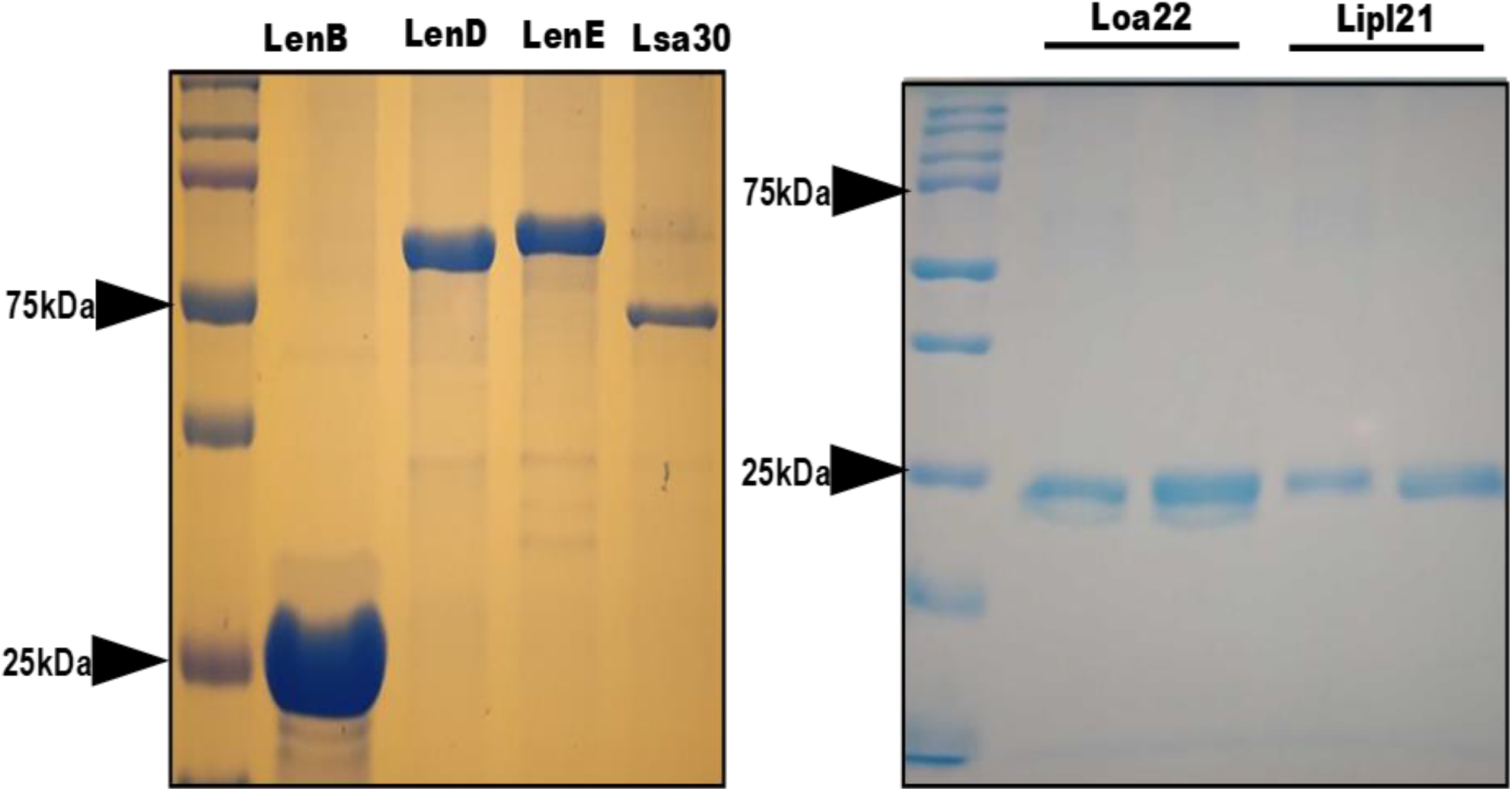
Purification of recombinant surface exposed lipoproteins: The recombinant proteins were purified as His-sumo fusion proteins as described in materials and methods. SDS PAGE profile shows the pure LenB (27kDa), LenD (61.94kDa), LenE (65KdA), Lsa30 (45kDa), Lipl21 (21kDa) and Loa22 (22kDa).

### Screening of recombinant proteins for their innate immune activity

To evaluate the innate immune activity of rLenB, rLenD, rLenE, rLsa30, rLipl21 and rLoa22 we tested their ability to activate macrophages. We stimulated mouse macrophages (RAW264.7), PMA differentiated human monocytes (THP-1), and bovine macrophages (BoMac) with varying doses (1, 2 and 5µg/ml) of the proteins. Our result shows that while Loa22 induced strong pro-inflammatory response in mouse, humans and bovine as evidenced by induction of IL-6 and TNF-α(Fig. 2 & Fig.3). Further these proteins induced concentration dependent pro-inflammatory response (Sup Fig. 1). Lsa30 induced very low or insignificant level of cytokines at concentration of 1µg/ml, however it induced significant levels of cytokines at higher concentration of 5µg/ml in all three host macrophages (Sup Fig. 1). LipL21 induced strong response in mouse macrophages, however the response was attenuated bovine and humans. To rule out that the observed pro-inflammatory effect was protein specific and not due to contaminating LPS, each protein was passed through Polymyxin B-agarose and were also pre-incubated with PMB in cell stimulation assays. 500ng/ml and 1 µg/ml LPS pre-incubated with PMB was used as control to check potency of PMB. The estimated concentration of LPS in final protein preparation varied from (0.10–0.15ng/ml)(Data not shown). The effect was protein specific because Proteinase-K plus heating abolished cytokine production. Besides, PMB inhibited the LPS induced cytokine production but did not attenuate the levels induced by proteins, indicating that the stimulatory effects observed were specific to protein and not due to contamination with LPS (Sup Fig. 2). To confirm whether stimulation with these proteins causes activation and maturation of macrophages we analysed the expression of costimulatory molecules (CD80, CD86) and maturation marker (MHC-II) in RAW2645.7 cells. All proteins induced expression of CD80, CD86 and MHCII, however maximum expression was induced by Loa22 followed by Len family of proteins (Fig. 2B). These markers were specifically induced by proteins as expression was significantly inhibited upon treatment with PK (Sup Fig. 3). Further PMB inhibited the LPS induced expression of these markers on mouse macrophages (Sup Fig. 3). These results indicate that the stimulatory effects induced by these proteins were not due to contamination with LPS.

**Fig.2:**
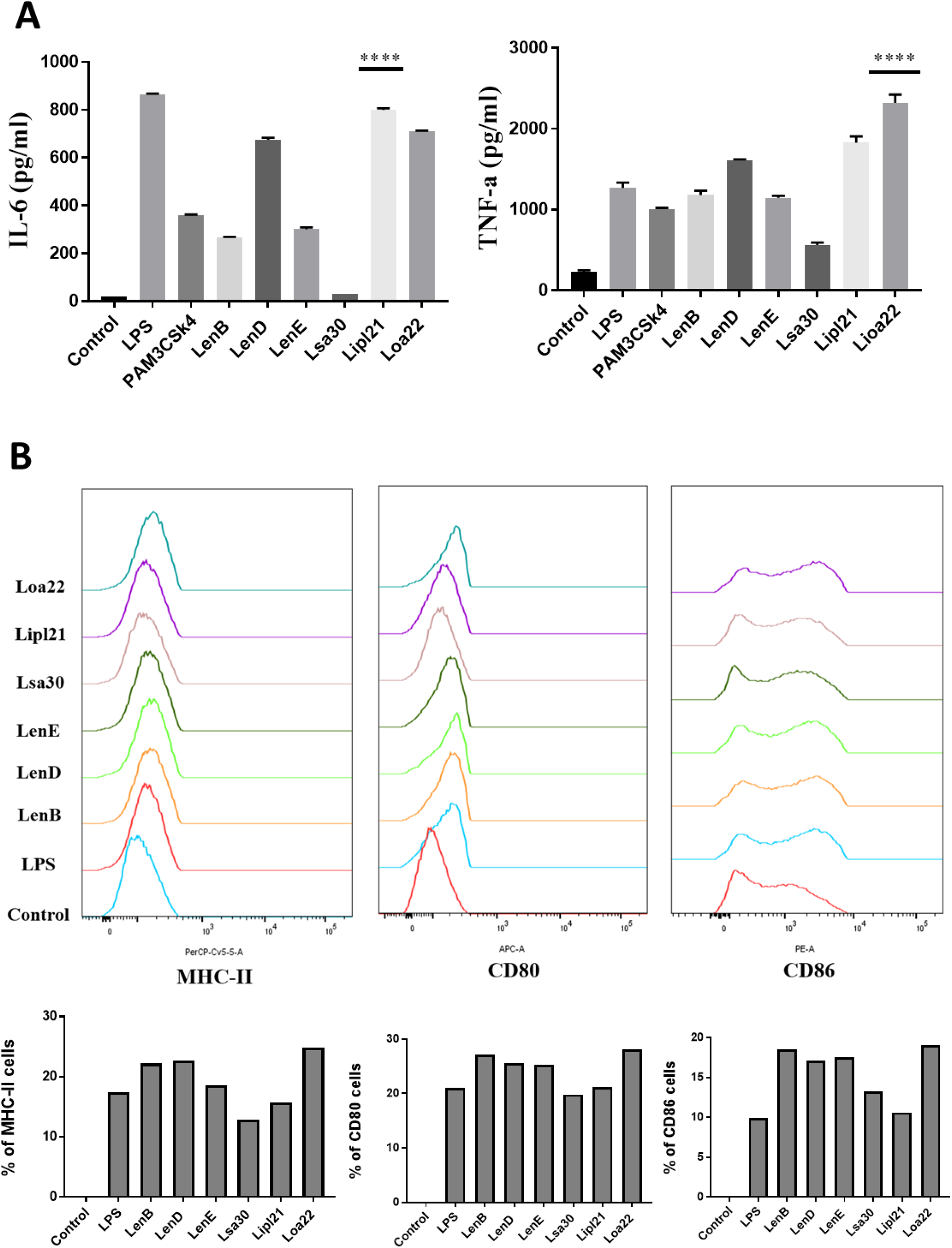
Analysis of activation of mouse macrophages after stimulation with surface proteins. (A) *Screening of pro-inflammatory response of surface proteins in RAW264*.*7 cells by ELISA*. RAW264.7 cell lines were stimulated with LPS (500ng/ml) or PMB treated 1ug/ml rLenB or rLenD or rLenE or rLsa30 or rLipl21 or rLoa22 for 24h and supernatant was collected to measure levels of TNF-α and IL-6 using sandwich ELISA kit. **(B)** *Expression of surface markers in RAW cells after stimulation with surface proteins*. RAW264.7 cell lines were stimulated with LPS (500ng/ml) or PMB treated 2ug/ml rLenB or rLenD or rLenE or rLsa30 or rLipl21 or rLoa22 for 24h (2ug/ml). Cells were stained with fluorochrome-conjugated antibodies against CD80, CD86, MHCII and then analysed by Flow cytometry. as described in materials and methods.

**Fig.3:**
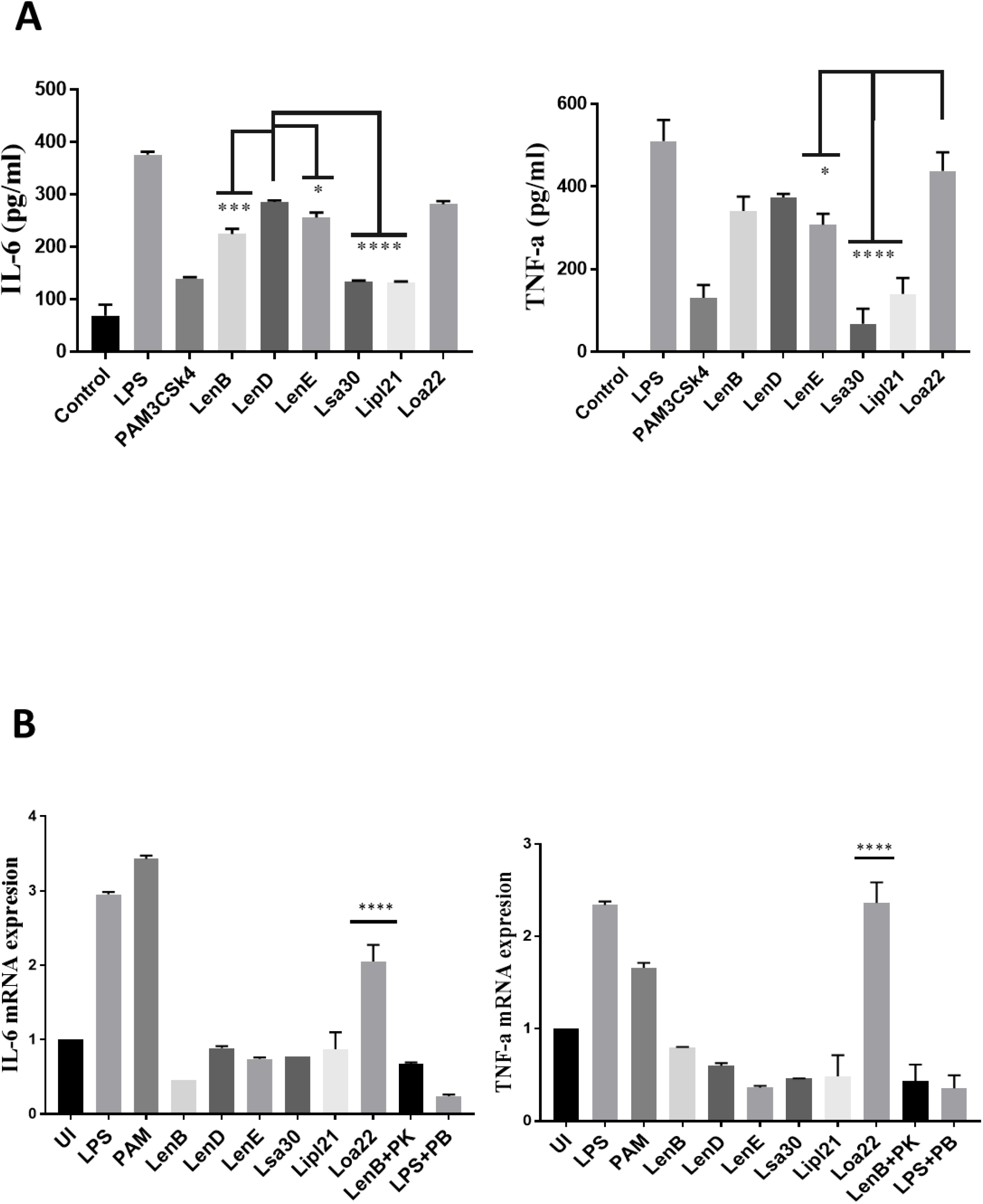
Proinflmmatory effects of surface proteins on human and bovine macrophages. (A) *Screening of pro-inflammatory response of surface proteins in THP-1 cells by ELISA*. PMA differentiated THP-1 cells were stimulated with LPS (500ng/ml) or PAM3CSK4(20ng/ml) or PMB treated 1μg/ml rLenB or rLenD or rLenE or rLsa30 or rLipl21 or rLoa22 for 24h and supernatant was collected to measure levels of TNF-α and IL-6 using sandwich ELISA kit. **(B)** *Screening of pro-inflammatory response of surface proteins in Bomac cells by RT-PCR*. BoMac cells were cultured and stimulated with proteins as described above. Cells were harvested, RNA isolated and converted cDNA. RT-PCR was performed using specific primers of bovine IL-6 and TNF-a as described in materials and methods (primer sequ.) Data are representative of three different experiments. Significant differences were calculated using ordinary one way ANOVA (****.***,**,* indicate P<0.0001,P<0.0002,P<0.0021,P<0.0332 respectively)

### Screening of surface proteins for their ability to bind to complement regulators (FH and C4BP) and host proteases (Plasminogen)

Previous studies have shown that several surface proteins including Len B, Len D Len E, Lsa30 and Loa22 of *Leptospira* bind to complement regulators like FH or C4BP. However, binding of these proteins was tested with any one of the complement regular either FH or C4BP and binding of second most abundant protein, LipL21 has not been tested yet. In this study we screened these proteins (Len B, Len D Len E, Lsa30, LipL21 and Loa22) for binding with both FH and C4BP. We also assessed their ability to bind host proteases (plasminogen, PLG) and mediate subsequent plasmin activity. Our dot blot result shows that all the proteins including LipL21 was able to bind to FH and C4BP (Fig. 4). Variable region of LigA (LAV) which was used as positive control showed binding with both FH and C4BP whereas BSA which was used as negative control didn’t show any significant level of binding (Fig. 4). This was further confirmed by ELISA where binding was enhanced with increasing concentration of NHS (0-100%, Sup Fig. 4). Further all proteins showed binding with PLG which was enhanced with increase in concentration of NHS (Fig. 5, Sup Fig. 4). The binding of proteins to PLG correlated to plasmin activity which was enhanced with time (Fig. 5B, Sup Fig. 5). Among the proteins tested, Loa22 demonstrated strongest binding affinity to PLG correlating to generation of significantly higher levels of plasmin (Fig. 5B, Sup Fig. 5).

**Fig.4:**
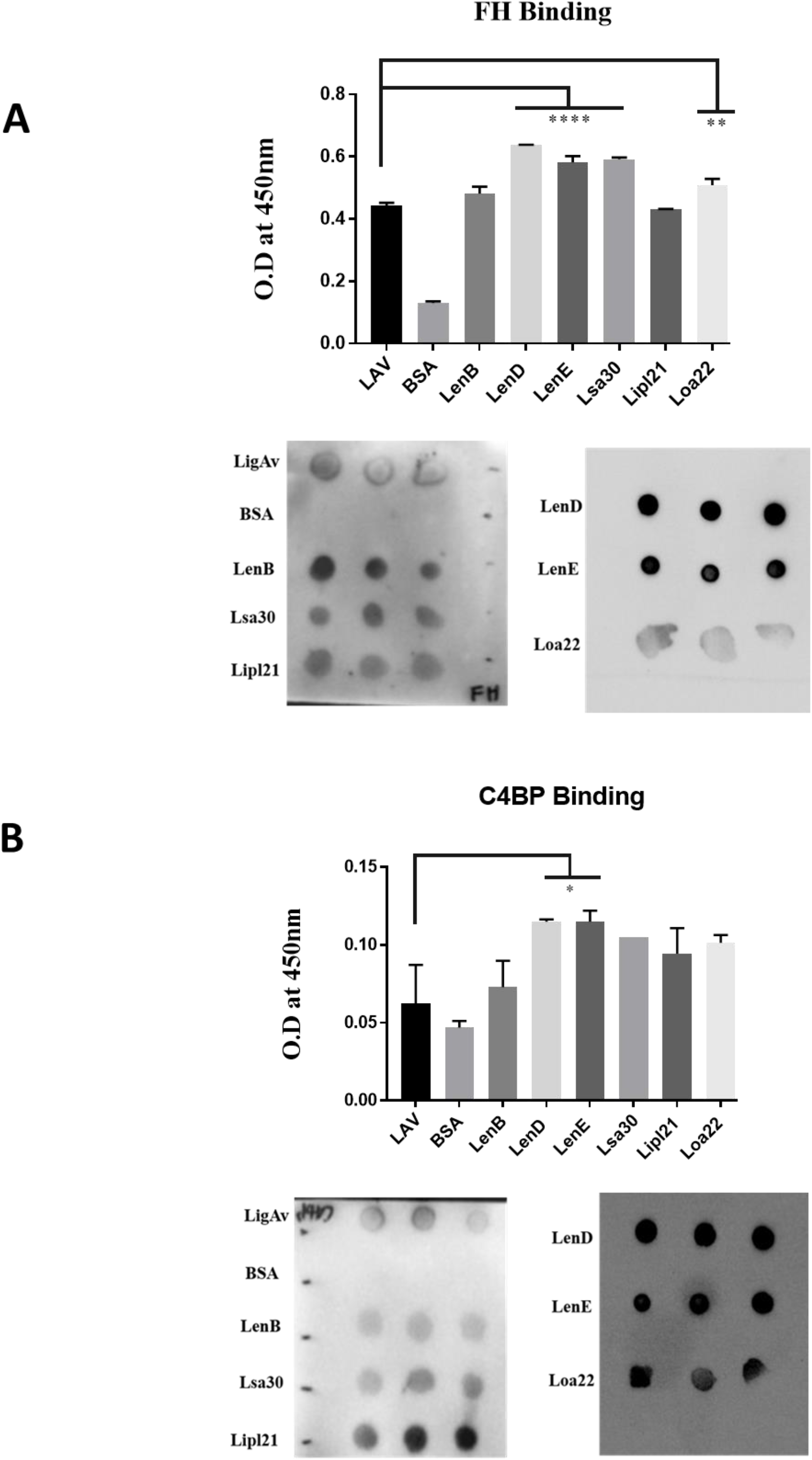
Evaluation of binding of surface proteins with complement regulators (FH and C4BP). (A) *Binding of surface proteins with FH and C4BP as analysed by Dot blot and ELISA*. Purified proteins, BSA (negative control), LAV (positive control), 1ug rLenB or rLenD or rLenE or rLsa30 or rLipl21 or rLoa22 were immobilized on nitrocellulose membranes and then incubated with 10%.. FH and C4BP were detected with specific antibodies by Western blot. (B) Microtitre plates were coated with 1µg/ml of proteins and binding was detected with specific antibodies against FH and C4BP as described in materials and methods. All data are representative of three independent experiments. Significant differences were calculated using the ordinary one way ANOVA (****.***,**,* indicate P<0.0001,P<0.0002,P<0.0021,P<0.0332 respectively)

**Fig.5:**
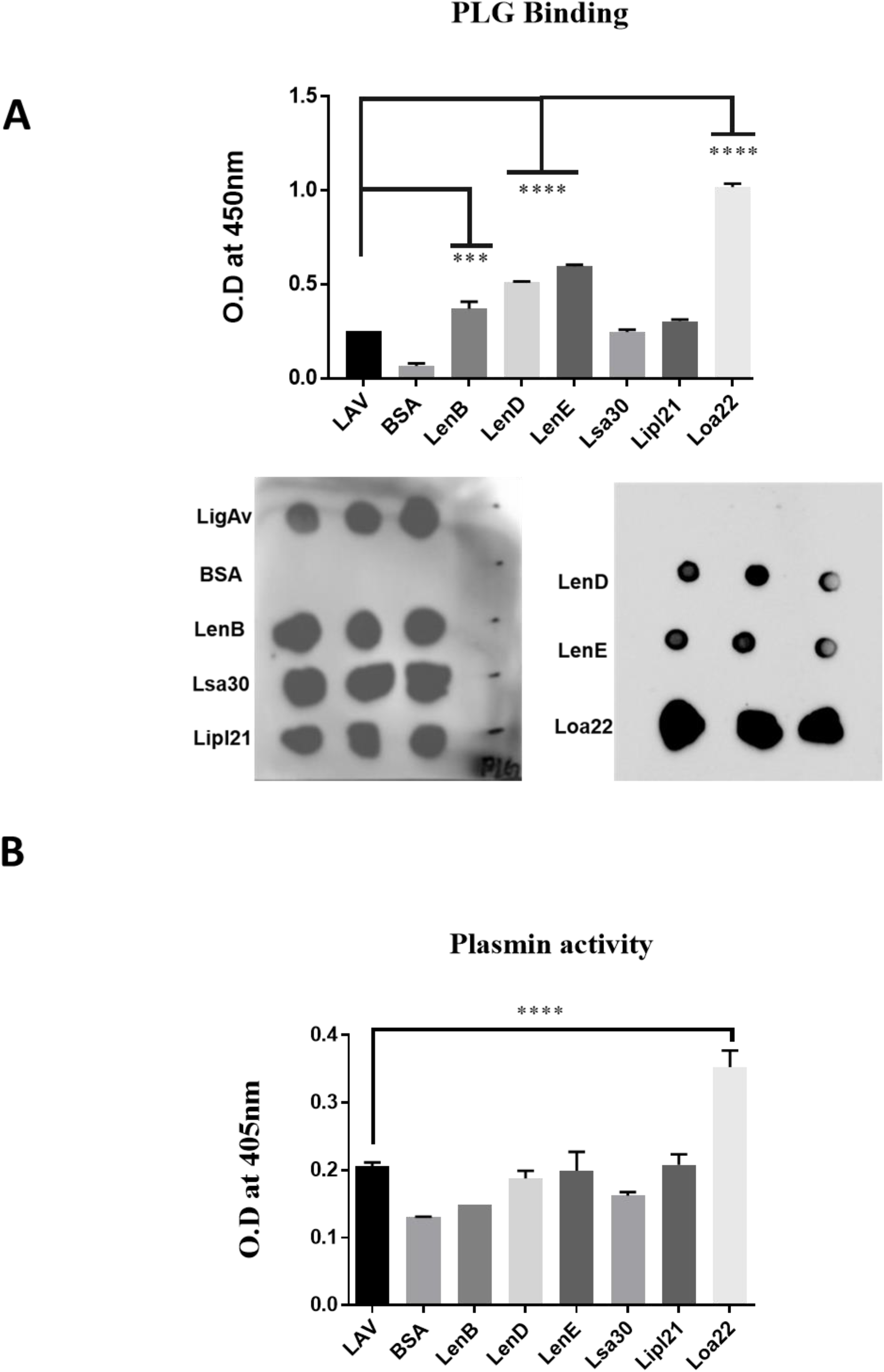
Evaluation of binding of surface proteins with host protease Plasminogen (PLG) (A) *Binding of surface proteins with Plasminogen (PLG) as analysed by Dot blot and ELISA* Purified proteins, BSA (negative control), LAV (positive control),), rLenB or rLenD or rLenE or rLsa30 or rLipl21 or rLoa22 were immobilized on nitrocellulose membranes and then incubated with 10% NHS (as a source of PLG). PLG were detected with specific antibodies by Western blot. Microtitre plates were coated with 1µg/ml of proteins and binding was detected with specific antibodies against PLG as described in materials and methods **(B)** *Plasmin activity*. Surface proteins (1ug/well) or BSA (1ug) were immobilized on microtiter plates followed by the addition of PLG, uPA and specific plasmin substrate. The plate was incubated for 72h and absorbance was read at 405nm as described in materials and methods. All data are representative of three independent experiments. Significant differences were calculated using the ordinary one way ANOVA (****.***,**,* indicate P<0.0001,P<0.0002,P<0.0021,P<0.0332 respectively)

### Screening of surface proteins for nuclease activity to evaluate their potential role in evasion from Neutrophil extracellular traps (NETs)

Several bacteria express nucleases to escape from Neutrophil extracellular trap (NET). *Leptospira* is known to induce NET, however whether they express nuclease to degrade NET is not known. In order to identify such nucleases, we screened nuclease activity of surface exposed lipoproteins Len B, Len D, Len E, Lsa30, LipL21 and Loa22 by incubating the DNA fragment with each protein (5µg) along with different metal ions (divalent cations) like Ca2+, Zn2+, Mg2+. Cleavage activity was quantified by measuring the ethidium bromide signal in each lane and calculating the fraction of DNA digested relative to the untreated DNA. Our result shows that only LenB, Lsa30 and LipL21 exhibited significant nuclease activity in presence of Ca2+, whereas all the proteins expect Loa22 exhibited nuclease activity in presence of Zn2+ (Fig. 6). LenB and partially LenE exhibited nuclease activity in presence of Mg2+. LenD and LenE showed strong activity in presence of Zn2+ whereas weak activity in presence of Ca2+ and Mg2+ (Fig. 6). LenB showed nuclease activity in presence of all metal ions whereas Loa22 didn’t show any nuclease activity irrespective of metal ions used (Fig. 6). To correlate the nuclease activity with *Leptospira* pathogenesis we tested the ability of these proteins to degrade PMA induced NET in vitro. Our result shows that all the proteins including Loa22 were able to degrade NET as evidenced by increase in levels of elastase due its release after treatment with protein (Fig. 7). The elastase levels were enhanced due to treatment with DNase which was significantly reduced when DNAse were pre-treated with EDTA (a chelating agent) indicating the inhibition of nuclease activity(Fig. 7).

**Fig.6:**
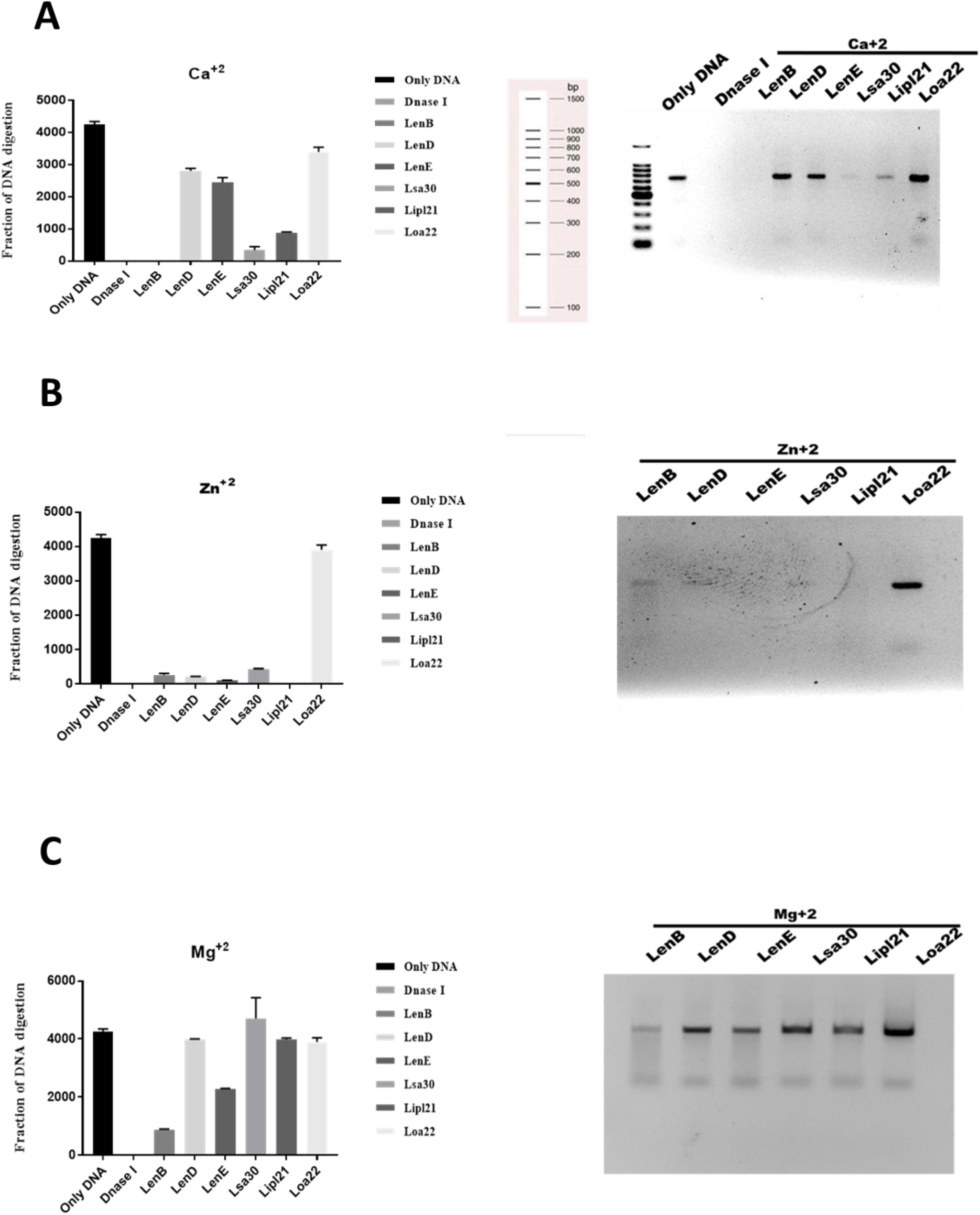
Evaluation of in vitro nuclease activity of surface exposed lipoproteins. *DNase activity of the surface proteins in presence of metal ions*. 700bp DNA fragment (200ng) was incubated with 5ug of surface proteins or DNaseI in DPBS with 5mM MgCl_2_ or CaCl_2_ or ZnCl_2_ at 37°C for 3h followed by visualization using the Agarose gel electrophoresis.

**Fig.7:**
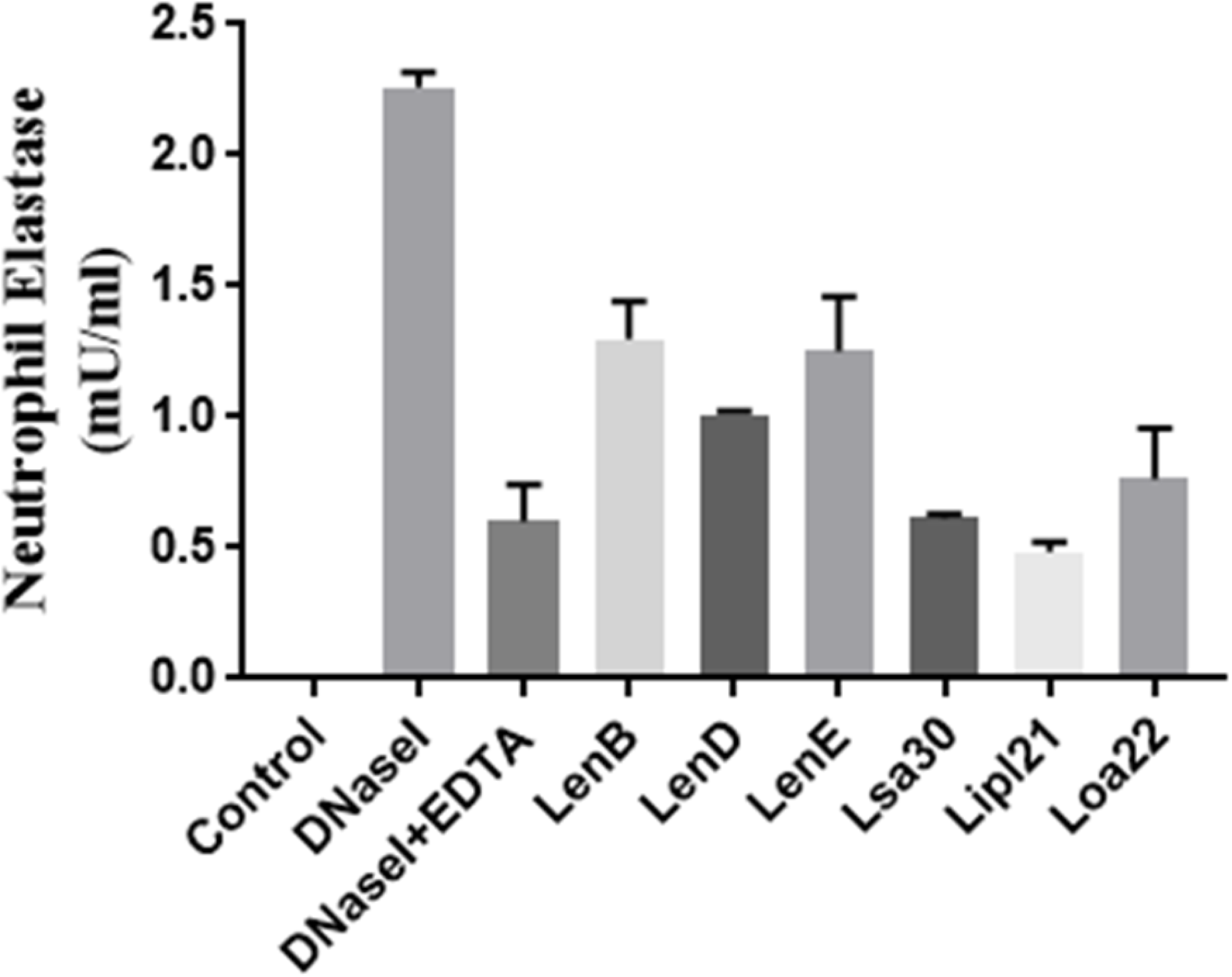
Evaluation of surface proteins ability to degrade Neutrophil Extracellular Trap (NET). Purified Neutrophil from mouse blood were stimulated with PMA to induce NET, followed by incubation with 5ug protein each protein in nuclease assay buffer for 4 hrs at 37°C /5%CO2 as described in materials and Methods. Elastase activity was determined by using Neutrophil elastase activity kit () using following manufacturers instruction. Data is representative of three independent experiments. Significant differences were calculated using the ordinary one way ANOVA (****.***,**,* indicate P<0.0001,P<0.0002,P<0.0021,P<0.0332 respectively)

## Discussion

Surface exposed lipoproteins of *Leptospira* play significant role in facilitating its survival in different environmental conditions and hosts for prolong periods (Haake, 2000). Several surface exposed lipoproteins of *Leptospira* have been either not characterized or are of unknown function (Raja and Natarajaseenivasan, 2015). The bacteria exploit these proteins to circumvent host mediated killing at various stages of infection. These proteins may act as adhesin and bind to host extra cellular matrix molecules like fibronectin, laminin, collagen etc. thereby helping bacterium to attach to host cell to initiate infection ((Christodoulides et al., 2017)). They may activate the innate response via signalling through toll like receptors like TLR2 or TLR4, inducing a pro-inflammatory response which may favour bacterial pathogenesis (Yang 2007, Yang et al. 2006, Faisal et al. 2016, Wang et al. 2012). Alternatively, to evade this innate recognition *Leptospira* like other bacterial pathogen may downregulate the expression or cause antigenic variations in their surface molecules including proteins upon infection in host(Cox et al., 1992;Zhang and Norris, 1998;Bunikis and Barbour, 1999;Xue et al., 2010). Further, the bacteria may utilize these proteins to evade from complement attack by binding to complement regulators like Factor H, C4BP or acquiring host proteases like plasminogen which cleaves complement component(Verma et al., 2006;Stevenson et al., 2007;Castiblanco-Valencia et al., 2012;Souza et al., 2012;Vieira et al., 2012;Vieira et al., 2014;Fraga et al., 2016).*Leptospira* like other bacterial pathogens may exploit these proteins to modulate function of phagocytes like neutrophils by inducing apoptosis, inhibiting phagocytosis and production of ROS or degrading Neutrophil extracellular traps (Kinkead et al., 2018;Kobayashi et al., 2018;Vieira et al., 2018;Ledo et al., 2020). Several surface proteins of *Leptospira* have shown binding to FH, C4BP and PLG and mediate co-factor activity thereby indicating their possible role in evasion from complement mediated killing(Verma et al., 2006;Stevenson et al., 2007;Barbosa and Isaac, 2020) (Castiblanco-Valencia et al., 2012;Souza et al., 2012;Wolff et al., 2013;Breda et al., 2015;da Silva et al., 2015;Fraga et al., 2016;Lin et al., 2016;Siqueira et al., 2016a;Siqueira et al., 2016b;Amamura et al., 2017;Salazar et al., 2017;Kochi et al., 2019). (Verma et al., 2006;Stevenson et al., 2007;Barbosa and Isaac, 2020) (Castiblanco-Valencia et al., 2012;Souza et al., 2012;Wolff et al., 2013;Breda et al., 2015;da Silva et al., 2015;Fraga et al., 2016;Lin et al., 2016;Siqueira et al., 2016a;Siqueira et al., 2016b;Amamura et al., 2017;Salazar et al., 2017;Kochi et al., 2019). Till now only LipL21 is known to modulate neutrophil function by inhibiting Myeloperoxidase (MPO) activity(Vieira et al., 2018). As reported in other bacterial species, surface proteins as nucleases which are involved in degradation of NET has not been reported in *Leptospira*.(Kiedrowski et al., 2014;Morita et al., 2014;Dang et al., 2016;Binnenkade et al., 2018). In spite of identification and characterization of several surface proteins of *Leptospira* involved in innate activation or complement evasion, not much effort is done towards identification of immunomodulatory surface proteins which are simultaneously involved in both immune activation and evasion. Recently our group has demonstrated the immunomodulatory role of *Leptospira* immunoglobulin like protein A and characterized the domain involved in activation and evasion from host innate immune response (manuscript in communication). Considering the functional redundancy in Leptospira surface proteins, in the present study we aimed to identify and characterize the surface exposed lipoproteins having this immunomodulatory or dual role that may help in better understanding their critical role in host pathogen interaction. Based on review of previously published studies we selected few proteins from *Leptospira* endostatin like protein family (LenB, LenD, Len E), *Leptospira* surface adhesin (Lsa30), 21kd Lipoprotein (LipL21), OmpA like protein (Loa22) that are known be involved in pathogenesis, and then deciphered their role in immunomodulation in terms of their ability to activate innate immune cells or evade from complement system or phagocytes(Raja and Natarajaseenivasan, 2015).

Our result shows that of various proteins tested Loa22 induced strong activation of macrophages of all hosts (mouse, human and bovine) while Lsa30 was least stimulatory in terms of cytokine production (Fig. 2, sup Fig.1). Further these proteins were able enhance the expression of costimulatory molecules (CD80, CD86) and maturation marker (MHCII) which is indicative of their ability to activate these cells (Fig 2). Previous studies have shown pro-inflammatory effects of surface proteins of *Leptospira* mainly on mouse macrophages, however, the ability of these proteins to activate macrophages of susceptible hosts (human, bovine) has not been tested yet (Wang et al., 2012;Faisal et al., 2016;Hsu et al., 2021). In our study we compared the innate activity of these proteins in macrophages of susceptible hosts (human and bovine) and our result shows that the stimulatory effects of these proteins varied in different host macrophages. Further the response against LipL21 in Human THP-1 and BoMac cells was attenuated as revealed by production of significantly lower levels of pro-inflammatory cytokines (Fig. 3). One can argue that the pro-inflammatory effect of these recombinant proteins might be due to low amount of contaminating LPS, however use of proper controls like Polymixin B (PMB) and Proteinase K (PK) in our assay has ruled out this possibility and gives us confidence that the observed stimulatory effect is attributed to inherent property of these proteins (sup. Fig. 2 &3). The activation of macrophages by these surface exposed lipoproteins might be TLR dependent, possibly through TLR2 as they are lipoproteins, however this needs to be confirmed by additional experiments. Previous studies have shown that several recombinant surface proteins from both *Leptospira* and other bacterial pathogens have shown to activate macrophages via signalling through TLR2 or TLR4 (Byun et al., 2012a;Byun et al., 2012b;Byun et al., 2012c;Sjolinder et al., 2012;Wang et al., 2012;Choi et al., 2016;Faisal et al., 2016). Loa22 has been shown to induce pro-inflammatory response via signalling through TLR2(Hsu et al., 2021) whereas LipL21 by virtue of its binding to peptidoglycan has been shown to be involved in immune evasion by escaping recognition from NLRs (Ratet et al., 2017). Ongoing work in our lab is focussed on identifying the host receptor involved in recognizing these lipoproteins leading to subsequent activation of innate immune cells like macrophages and DCs.

*Leptospira* is known to evade complement attack via binding complement regulators, secretion of self-proteases or acquiring host proteases to degrade complement components(Amamura et al., 2017). Several surface exposed lipoproteins of *Leptospira* apart from binding to components of host extracellular matrix have also been reported to bind to complement regulators and host proteases like plasminogen(PLG)(Siqueira et al., 2016a;Siqueira et al., 2016b;Kochi et al., 2019). Previous studies have shown binding of Len A and Len B to FH and Lsa30 to C4BP and PLG(Stevenson et al., 2007;Souza et al., 2012). Binding of other members of Len family (Len C, LenD, LenE and Len F), LipL21 and Loa22 with complement regulators or PLG has not been evaluated yet. Our study not only confirms the previous reports but also demonstrates the binding of these proteins to both FH and C4BP and also to host proteases PLG, thereby highlighting their possible role in evading complement mediated killing via inhibiting both classical and alternate pathways (Fig. 4 &Fig. 5).

Phagocytes like Neutrophils have evolved a novel mechanism of killing the bacteria in extracellular space by throwing their DNA, along with histones and antimicrobial peptides which are referred as Neutrophils extracellular traps (NETs)(Brinkmann et al., 2004). This mechanism of killing extracellular bacteria by trapping outside the cell is called NETosis and is independent of phagocytosis and degranulation(McDonald et al., 2012). However, pathogens have evolved strategies to suppress or degrade NETs by expressing surface proteins having nuclease activity(Kaplan and Radic, 2012). *Leptospira* upon encounter with neutrophils can induce NET, however to evade from NETosis whether it expresses nucleases has not been reported yet(Scharrig et al., 2015). Our group for the first time reported the role of *Leptospira* immunoglobulin like protein A (LigA) in degrading NET by virtue of its nuclease activity (manuscript in communication). Considering the functional redundancy in surface proteins of *Leptospira* it is likely that nuclease activity might also be possessed by other surface exposed lipoproteins. Keeping this in view we tested the nuclease activity of Len B, Len D Len E, Lsa30, LipL21 and Loa22. Our result showed that all the proteins except Loa22 exhibited nuclease activity in vitro albeit with requirement of different metal ions (Fig. 6). Further, the nuclease activity of these proteins correlated to their ability in degrading NET in vitro and thus highlights their possible role in escaping the *Leptospira* from NETosis (Fig. 7). Although Loa22 didn’t showed any nuclease activity in vitro, however it was able to degrade NET as evident from elastase activity (Fig. 7). We speculate that Loa22 apart from having exonuclease activity might also possess endonuclease activity which needs further investigation. LipL21 can modulate neutrophil function by inhibiting Myeloperoxidase (MPO) hence it is likely that apart from nuclease activity these proteins might have role in modulating neutrophil function by inhibiting ROS, MPO, phagocytosis or inducing apoptosis. However these aspects needs to be tested.

In conclusion, our results demonstrate that surface exposed lipoproteins of *Leptospira* tested in this study have role in modulating the host innate response by inducing pro-inflammatory effect, bind to complement regulators and host proteases to evade from complement attack, degrade NET by virtue of nuclease activity. Thus, identification and characterization of immunomodulatory surface exposed lipoproteins is of significant importance in understanding *Leptospira* pathogenesis. Further, deciphering the immunomodulatory or dual role (immune activation and evasion) of these proteins may also help in understanding how the bacteria disseminate and colonize in various organs which may provide important insight into clinical outcome in various hosts.

## Acknowledgements

This work was supported partly from DBT project-BT/PR21430/ADV/90/246/2016 (SP025) on developing *Leptospira* vaccines and partly from DBT-NIAB flagship project No-BT/AAQ/01/NIAB-Flagship/2019 (SP051) on host-pathogen interaction which are funded to SMF from the Department of Biotechnology, Ministry of Science and Technology, Government of India. Financial support from the NIAB core fund is duly acknowledged. The authors would like to thank the Director, NIAB, Dr. Subeer S. Majumdar for providing necessary infrastructural facility and support for the execution of the above study. AK is supported by UGC fellowship and registered for PhD programme at RCB, Faridabad. VPV is supported by CSIR fellowship and registered for PhD programme at Manipal University, Manipal.

## Author’s contribution

SMF conceived the idea and designed the experiments. AK and VPV performed the experiments. AK, VPV and SMF analyzed the data. AK and SMF wrote the initial draft and SMF edited the manuscript. All authors approved the final version of the manuscript.

